# Biocontrol of PGPR strain *Bacillus amyloliquefaciens* Ba168 against *Phytophthora nicotianae* on tobacco

**DOI:** 10.1101/700757

**Authors:** Dongsheng Guo, Chenhong Yuan, Yunyan Luo, YaHan Chen, Meihuan Lu, Guochan Chen, Guangwei Ren, Chuanbin Cui, Jiatao Zhang, Derong An

## Abstract

Tobacco black shank (TBS) caused by *Phytophthora nicotianae* is destructive to almost all kinds of tobacco cultivars and is widespread in many tobacco-planted countries. Here, an isolated plant growth-promoting rhizobacteria (PGPR) strain Ba168 is promise in biocontrol of TBS. *In vitro* assays disclosed a strong *P. nicotianae* suppression activity and the field utilization potential (FUP) by characterized the crude extract of culture filtrates of Ba168. *P. nicotianae*’s growth was inhibited by the crude extract at minimum inhibitory concentration (MIC) of 5μl/mL. Extracellular conductivity, pH and the wet, dry weight of *P. nicotianae*’s mycelia, were significantly different after treated with different concentrations of the crude extract and the deformity and perforation of treated *P. nicotianae*’s hyphae can be observed in scanning electron microscope (SEM) analysis. Proteome characterizations of the crude extract were used as supplementary proofs that further evaluated FUP of Ba168. We then identified strain Ba168 as *B. amyloliquefaciens* by its genetic and phenotypic characteristics. Field assays comparatively evaluated TBS control efficacy of these PGPRs and agrochemicals. Pooling analysis of the results showed that the biocontrol efficacy of Ba168 preparation is only lower than Mixture of Propamocarb hydrochloride and Azoxystrobin (MPA) but better than other tested subjects. Although the existence of differences in biocontrol efficacy, PGPR preparations effectively reduced the disease index of tobacco.

**Importance:** This work demonstrates the promising biocontrol potential of B. amyloliquefaciens Ba168 and highlights the positive roles of PGPR in suppression of this soil-borne disease.

---

TBS is a kind of soil-borne disease caused by *P. nicotianae* that damages the leaf, stem and root of tobacco in the whole growth period of the tobacco plant (1). This pathogen would spread rapidly under conditions of high temperature and high humidity, causing high yield loss (2–4). The traditional countermeasures of crop rotation, breeding for resistant tobacco varieties and application of pesticides are insufficient to control this soil-borne disease (5). Compared to crop rotation, counterintuitively, monoculture systems like planting the same crop only every 4-5 years by using pathogen-free seeds will to some extent prevent losses owning to soil-borne pathogen like *Gaeumannomyces graminis* var. *tritici.* However, as it is not practical for economic reasons, these pathogens commonly have devastating effects on greenhouse and field (5). Pesticides are the most commonly used tools, sometimes effective, but the widespread use of it over a long term will cause severe environmental problems and, more importantly, develop resistant strains (6–8).

Biocontrol of soil-borne pathogens is preferred worldwide, because of the favorable biocontrol efficacy, if applied properly, and the urgent need to introduction environmental friendly alternatives to gradually reducing the use of harmful agrochemicals (9). So far, various *P. nicotianae* control strains, like *Bacillus subtilis* Tpb55 (10), *Bacillus atrophaeus* HAB-5 (11), non-pathogenic binucleate *Rhizoctonia* fungi (12), *Trichoderma* (13, 14), *Brassica carinata* pellets (15), *Glomus mosseae* and *Pseudomonas fluorescens* (16), have been evaluated. Among these biocontrol agents, *P. fluorescens* is one of the most studied PGPR (17), which contributed greatly to the understanding of the mechanisms that are involved in disease suppression. But the relatively unstable biocontrol efficacy, shorter storage period and toxic cyanides produced by it (18), to some extent, limited its application in the field recent years. At present, *Bacillus* products are gradually gobbling up the market share of *P. fluorescens*, because of the ability to inducing systemic resistant (ISR) responses (19), secreting diverse antimicrobial peptide and enzymes, which are harmless to humans, animals and plants (20, 21), improving plant growth by promoting absorption of nutrients, increasing nutrient availability, helping plants to adapt to a number of environmental stresses and improving the nutritional status (22). More importantly, spores produced in their natural state are resistant to stress and the size of spores perfectly satisfy the requirements of the production of preparations (9, 23). Biocontrol PGPR, sometimes irrespective of antibiotics production, elicit ISR, which allows plants to withstand pathogen attack to leaves or roots (24, 25). Bacillus-mediated ISR in plant disease suppression has been well studied in many cases (19, 26, 27). It also works in *P. nicotianae* suppression. A related study conducted by Wu et al show that root colonization by *B. amyloliquefaciens* FZB42 would elicit ISR and restrict stomata-reopening mediated by the *oomycete* foliar pathogen *P. nicotianae* in *Nicotiana benthamiana* and thus block this pathogen to initiate penetration and infection to some degree (28). But barely have studies focused on whether *Bacillus* works in *P. nicotianae* suppression in field. Besides, Characteristics of extracellular metabolites of a PGPR to some extent represent the biocontrol potential of it (29).

In this study, we (i) revealed biocontrol potential of Ba168 by evaluated *P. nicotianae* suppression activity of Ba168, (ii) further reflected FUP of Ba168 by analyzing the proteins in its culture filtrates, (iii) characterized strain Ba168 through morphological and molecular biological tools, (iv) evaluated TBS control efficacy of Ba168 and 5 PGPR preparations and 2 pesticides in 2 tobacco varieties over 2-year field assays. This work show us the effect of the extracellular metabolites of *B. amyloliquefaciens* directed at *P. nicotianae,* more importantly, vindicate PGPR’s positive effect in field and provide us an effective TBS control PGPR, *B. amyloliquefaciens* Ba168.

## Materials and methods

### Materials

*Alternaria alternata, Botrytis cinerea, Ralstonia solanacearum, P. nicotianae* were isolated from various *Nicotiana tabacum* that planted in Shaanxi Province and preserved in Plant Virus and Microorganism Resource Laboratory in the College of Plant Protection, Northwest A&F University.

1 newly emerged pesticides, MPA (Lier Chemical Co., Ltd., China), and 1 commonly used Chemical fungicides, 80% Dimethomorph water dispersible granules (Jiang su hui feng Co. LTD, China), were purchased from local market.

Biocontrol strains were all isolated from soil samples collected from intact forest of Qinling, Mountains, China. *Bacillus licheniformis* (isolated from deciduous broad-leaved forest in 2008), *B. subtilis* (isolated from bushes in 2000), *Brevibacillus laterosporus* (isolated from coniferous forest in 2009), *B. methylotrophicus* (isolated from deciduous broad-leaved forest in 2011)(30) and *Bacillus pumilus* (isolated from coniferous forest in 2001) were preserved in Plant Virus and Microorganism Resource Laboratory in the College of Plant Protection, Northwest A&F University. Strain Ba168 (isolated from evergreen broad-leaved forest in 2014) was deposited as strain 6426 in China General Microbiological Culture Collection Center (CGMCC).

Tobacco varieties used in field trials were *N. tabacum* QinYan96 and NC89 (provided by Shaanxi Provincial Tobacco Research Institute).

### Preparation of *P. nicotianae* spores, Ba168 crude extract and field trial agents

Spore suspensions of *P. nicotianae* were prepared according to the method described by Kong et al(10). Finally, the spore suspension was adjusted to 10^6^ CFU/mL by hemocytometer.

Agrochemicals involved in field trials were diluted at a ratio of 1/2500 of the original concentration.

6 biocontrol agents containing spores and secondary metabolisms, were identically prepared. Strains were inoculated on Luria-Bertani (LB) plates and then cultured at 27 °C for 24 h. Further, a single colony was selected and inoculated in conical flasks containing LB broth (150 mL conical flasks, 100 mL per each) and cultured in a shaking table at 150 r/min, 30 °C for 48 h. After that, the fermentation broth was inoculated in a sterile solid fermentation medium in a volume ratio of 1/10. For instance, about 1 kg powder can be generated by fermentation raw material which was composed of sucrose 100 g, yeast extract 100 g, MnSO_4_ 0.2 g, MgSO_4_ 1 g, soybean meal 600 g, bran 600 g, water 1800 mL. After preparation, the raw material was packed in a 250 mL conical flask(80 g per flask) and incubated at 28 °C for 72 h. Finally, the semi-processed raw material was dried in a 40-degree oven for 96 h, and then crushed by a grinder, and the powder which across a 200-mesh sieve was collected. More spores can be generated if we using this collected powder as inoculum (1g/ 10ml sterile water) to replace fermentation broth.

The crude extract of extracellular metabolites of strain Ba168 was prepared by the following steps. Medium used in this part is Inorganic nitrogen source medium (INSM) composed of 20 g/L glucose, 20 g/L (NH_4_)_2_SO4, 1 g/L K_2_HPO_4_, 0.5 g/L MgSO_4_, 0.5 g/L NaCl. The pH should be adjusted to7.0. Initially, Strain Ba168 was inoculated to the 250 mL conical flask containing 150 mL INSM medium and cultured in a shaking table at 150 r/min, 28 °C for 24 h. The fermented liquid subsequently separated in a 50 mL centrifuge tube and centrifuged at 10000 r/min for 20 min. Removal of precipitation and join in different quality of ammonium sulfate in a 50mL centrifuge tube, and make the saturation of ammonium sulfate are 10%, 20%, 25%, 30%, 35%, 40%, 50%, 60%, 70%, 80%, 90%, 100%. Then we kept those tubes on a test-tube rack at a fridge (4 °C) for 24 h, afterward, centrifuged again, abandon supernatant, sediments in each tube were dissolved in phosphate buffer (25 mmol/L PH=7.4) of 0.5 mL, dialysis desalting. The crude extract of Ba168 was the combined final products across a bacteria filter (0.22 um, MILLEX).

### The inhibitory efficiency of 5 *Bacillus* strains compared to Ba168 on *P nicotianae*

Dual culture assay (31) was used to determine the antifungal activity of these strains. After the mycelia of *P. nicotianae* fully covered Potato Dextrose Agar (PDA) plate, using a 5 mm diameter sterile cork borer to harvest fungus samples along the colonies’ edge. Then, put the fungus cake at the center of a 90 mm wide petri dish (containing 10 mL PDA) and then 1μl overnight cultures (OD_600_ = 0.8) of each strain was aseptically pipetted at four perpendicular sites by a 2.5μl pipette, and each site was 25mm from the fungus block. After that, the petri dish was incubated at 27 °C for 7 days. During the interval of these days, we observed the antagonistic efficacy of each strain by calculating the average value of two perpendicular diameters of the inhibitor zone.

### The inhibitory efficiency of strain Ba168 on 4 tobacco pathogens

The inhibitory efficiency of 3 mycelial fungus *(A. alternata, B. cinerea, P. nicotianae*) was estimated by the method consistent with 1.3.1, whereas bacterial pathogen *R. solanacearum* was evaluated by using filter paper method (32). Filter paper discs (5 mm diameter) were submerged in antagonist suspensions. 100 μL diluted suspension (OD_600_ = 0.8) of *R. solanacearum* was spread inoculated onto LB agar in a Petri dish and allow air-dried for several minutes. Then, 4 discs were placed on the inoculated LB plates at four perpendicular sites.

### MIC of Ba168 crude extract

The MIC of the crude extract of Ba168 against *P. nicotianae* was determined following the method of Tian et al (33) with slight modifications. Initially, different volumes of crude extract of Ba168 was pipetted and dissolved separately into sterile water and adjust the volume to 0.5mL. We then melted PDA medium and cooled to 45-50 °C. After that, we aseptically pipetted PDA of 9.5mL onto 90mm wide glass Petri dishes and mixed uniformly with the 0.5 mL solution. Finally, the plates of the experimental group mixed with different concentrations (1, 2, 3, 4, 5, 10, 20, 50 uL/mL) of crude extract were prepared and set aside. Control plates (without Ba168 crude extract) were prepared in the same procedure. The disk of the mycelium of *P. nicotianae* harvested from the periphery of the four-day-old colony using a sterilized puncher (5mm in diameter), was inoculated at the center of each petri dish. The plates were incubated in an incubator with a temperature of 27 °C. All subjects were performed triplicate. The antifungal efficacy of each treatment was evaluated three days later and then recorded results daily for seven days, by observing the mean of two perpendicular diameters of each colony (34). The lowest concentration of crude extract that fully inhibited the growth of *P. nicotianae* was considered the MIC.

### Effect of crude extract of strain Ba168 on mycelial weights of *P. nicotianae*

Effect of crude extract of strain Ba168 on mycelia weights of *P. nicotianae* was evaluated by the method developed by Tian et al (33) and modified by Jing et al (35) with slight alterations that 1%(v/v) of spore suspension (4×10^6^ /mL) of *P. nicotianae* was inoculated into a 50 mL conical flask containing 20 mL Potato Dextrose Broth (PDB) which contain crude extract of strain Ba168 at 0, 1, 2, 3, and 4μl/mL, respectively. After incubation, the mycelia produced in liquid cultures were filtered by using Miracloth (Millipore) and washed three times by PBS. We weigh the wet weight of each group when droplets barely drop from Miracloth, whereas the dry weight of each mycelium was determined after drying at 60 °C for 24 h.

### Cell wall and cell Membrane Permeability Test

Experiments were conducted according to the methods of Paul et al (36) and Jing C. et al (35) with minor modifications. The 4×10^6^ /mL spore suspension of *P. nicotianae* was inoculated in a conical flask (250 mL) containing 150 mL OB(agar free oats medium) at an inoculation volume of 1% and incubated in a 150 r/min constant temperature shaker at 30 °C for 36 h. The mixtures were subsequently centrifuged at 3000×g/min for 10min. The precipitates then were collected and washed three times with sterilized H_2_O. After exposure the mycelium of *P. nicotianae* into the solution of crude extract in a concentration of 0, 1, 2, 3 or 4×μl/mL, we recorded extracellular conductivity by the electrical conductivity meter DDS-307 (Shanghai Electrical Instrument co., LTD., China) at 0, 30, 60, 90, 120min respectively, and extracellular pH by the pH meter FiveEasy Plus (METTLER TOLEDO., Switzerland) at 0, 15, 30, 45, 60, 75, 90, 105 min respectively.

### Effect of crude extract of strain Ba168 on the morphology of mycelia of *P. nicotianae*

The preparation of the mycelia of *P. nicotianae* following the method described in 1.5. The collected mycelia washed three times by PBS and then inoculated into a 50 mL sterilized centrifugal tube containing 40 mL phosphate buffer saline (PBS) with 2% (w/v) glucose and crude extract of different concentrations(0, 1, 4 ×μl/mL). Further, these tubes were incubated in a 150 r/min constant temperature shaker at a temperature of 27 °C for 12 h. Then we observed the collected mycelia under liquid nitrogen cooled Scan Electron Microscope (Nova Nano SEM 450, FEI, American).

### Proteome analysis of the crude extract of Ba168

An effective Biocontrol strains are capable of helping host plants withstand pathogens, like inducing resistant responses, producing various antibiotics and competing inches around the roots of host plants. On the other hand, strains should also have plant growth promoting abilities, like secreting phytase, auxin.

The analysis of all proteins in Ba168 culture filtrates may, to some extent, reflect the metabolites secretion of Ba168 after rhizosphere colonization in the field and further evaluate the potential application of Ba168 in the field combined with the experimental results described above.

For this reason, in this study, we proposed to use extracellular proteome analysis as a supplementary data to evaluate whether strain Ba168 have the potential to be utilized in field. We identified unique proteins related to biocontrol capacities of Ba168 by using liquid chromatograph-mass spectrometer (LC-MS) (Thermo Fisher) and molecular weight greater than 5000 can be identified. The crude extract without dialysis desalting was prepared.

### Identification of strain Ba168

The identification of strain Ba168 based on Gram staining, morphology, physiological and biochemical tests according to Bergey’s Manual of Determinative Bacteriology (37). Further identification of Ba168 was confirmed by 16SrRNA analysis. The forward primer (5’-AGTTTGATCMTGGCTCAG-3’) and the reverse primer (5’-GGTTACCTTGTTACGACTT-3’) was used (11). The phylogenetic analysis was conducted using representatives of *B. amyloliquefaciens* strain SB 3297 (Genbank accession number(GAN): GU191913.1) (38), *Bacillus vallismortis* strain DSM 11031 (GAN: NR_024696.1) (39), *B. subtilis* subsp. *Spizizenii* (GAN: AF074970.1) (40), *B. subtilis* strain IAM 12118 (GAN: NR_112116.2), *B. atrophaeus* strain ATCC 51189 (GAN: EF188847.1), *B. methylotrophicus* strain Y37 (GAN: KT890344.1) (41), *B. licheniformis* strain ATCC 14580 (GAN: NR074923.1) (42). *B. pumilus* strain ATCC 7061 (GAN: NR043242.1) (43) and *Bacillus cereus* ATCC 14579 (GAN: NC004722.1) (44). The reference strain *B. laterosporus* IAM 12465 (GAN: NR037005.1) (45) was used as an out-group.

The amplified procedure following the method described by Jangir et al (46) and the amplified RNA fragments was sequenced (Qingke, XiAn, China). The corresponding sequences of 16SrRNA were aligned by using MAFFT (https://mafft.cbrc.jp/alignment/server/index.html). Phylogenetic trees were constructed with MEGA 7.0 (47)software by using the Maximum Likelihood method with 1000 bootstrap replications.

### Field test of 6 biocontrol agents and 2 pesticides

The test sites was the places that TBS occurs frequently. Test sites A, B lies in the town of MaPing, XunYang county, AnKang city, Shaanxi Province. This location is 1020 meters above the sea level, with the longitude of 109 °08′ 15″ and latitude of 32°54′ 41″. From 2017 to 2018, tobacco varieties Qinyan96 were used in A, whereas NC89 was planted in B.

The test site A and B, was sandy loam with medium soil fertility, convenient for irrigation and drainage. Available phosphorus: 13.45 mg/kg, available potassium: 108.12 mg/kg, alkali-hydrolysis nitrogen: 52.45 mg/kg, total phosphorus: 0.93 g/kg, total nitrogen: 0.29 g/kg, organic matter content: 11.20 g/kg, pH=6.9.

2 test sites were identically prepared. There were 9 experimental groups and 1 control group in each site, and each group was performed four times. The spacing between rows and columns was 0.8 m and 1.1 m, and 100 (10×10) plants were planted in each plot. A single plot covers an area of 100 m^2^, and 20 rows of protection rows are planted on each side.

In experimental groups of biocontrol strain, tobacco plants was treated with seed treatment @5 g/kg + seedling dip @5 g/l before transplant in experiment sites, whereas pesticides and control groups without ferments treatment were transplanted simultaneously.

The first field application were on June 18 when the growth period of tobacco was the rosette stage. Mild symptoms of tobacco plants infected by *P. nicotianae* can be observed in that stage. Preparations was applied as a combination of + root inoculation @750 g/ha + foliar spray @ 750 g/ha. Before field application, solid ferments of biocontrol strains were diluted into suspensions @ 2 g/L. Root inoculation of preparations was performed by pipetting 30 ml of the bacterial or fungicide suspension onto the roots of tobacco plants. Foliar spray was complete by a manual sprayer. Control groups were root-inoculated and sprayed with the same dosage of sterile distilled water. Applying every 10 days for a total of 3 times.

Before the first treatment, the initial disease index was investigated, and the disease index was investigated 10 days after each treatment. Each plot was sampled at 5 points, the disease index of 10 tobacco plants were recorded at each point, and the total number of investigated tobacco plants and the number of diseased tobacco plants at each grade was recorded. The disease index was computed and shown below as previously described (10, 35, 48).

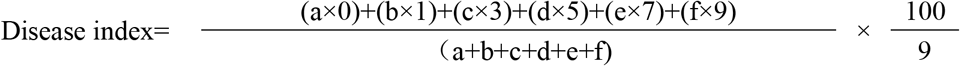

Where 0=no symptoms; 1, 3, 5, and 7=1/3, 1/3 to 1/2, 1/2 to 2/3, and 2/3 of the total leaves or the periphery of stems were wilted, respectively; 9 = tobacco plant was dead; a, b, c, d, e, and f note that the numbers of tobacco plants in each disease grade.

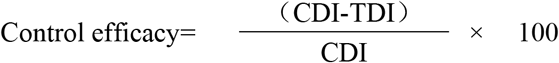

Where CDI refers to the increase of disease index of the control group, TDI means the rise of disease index of every treatment.

All data were analyzed using SPSS 22.0 with Duncan’s new multiple range test (DMRT) (P-value of <0.05) and were expressed as the mean ± standard deviation (mean ± SD) from three parallel experiments. The graph was built by Excel 2013 after data processing.

## Results

### Bicontrol potential test of strain Ba168

The Inhibitory Efficiency of 5 *Bacillus* strains compared to Ba168 on *P. nicotianae* was described in Table 1. The inhibition zone diameter of *P. nicotianae* (Fig 1) is highest, reached (28.42 ± 2.84). Moreover, Ba168 also showed good antagonistic activity against the other 3 tobacco pathogens including *A. alternate, B. cinerea* and *R. solanacearum* (Table 2). Antimicrobial activity of Ba168 showed in this test encourages further investigation.

**FIG 1.**
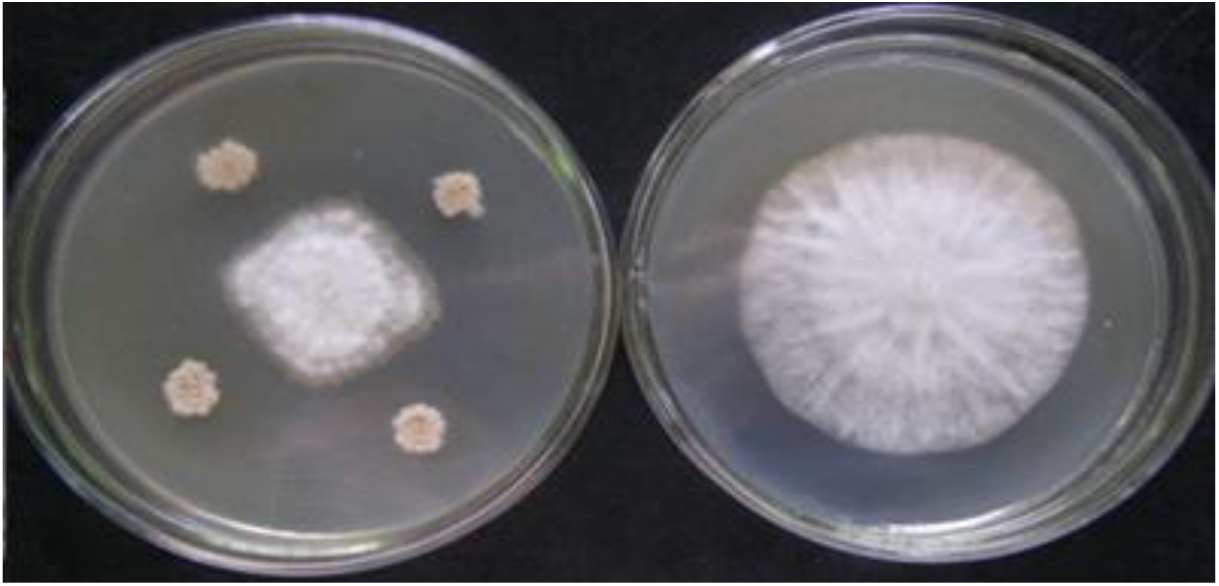
Dual cultures plate assay of strain Ba168 against *P. nicotianae*

**TABLE 1.** The inhibitory efficiency of 6 biocontrol strains

| Number | Pathogen | Inhibition zone diameter (Mean ± SE) |
| --- | --- | --- |
| 1 | strain Ba168 | 28.42 ± 2.84a |
| 2 | *B. licheniformis* | 22.55 ± 0.17c |
| 3 | *B. subtilis* | 25.88 ± 0.73b |
| 4 | *B. laterosporus* | 24.70 ± 0.93b |
| 5 | *B. methylotrophicus* | 26.38 ± 2.95b |
| 6 | *B. pumilus* | 22.01 ± 0.42c |

**TABLE 2.**
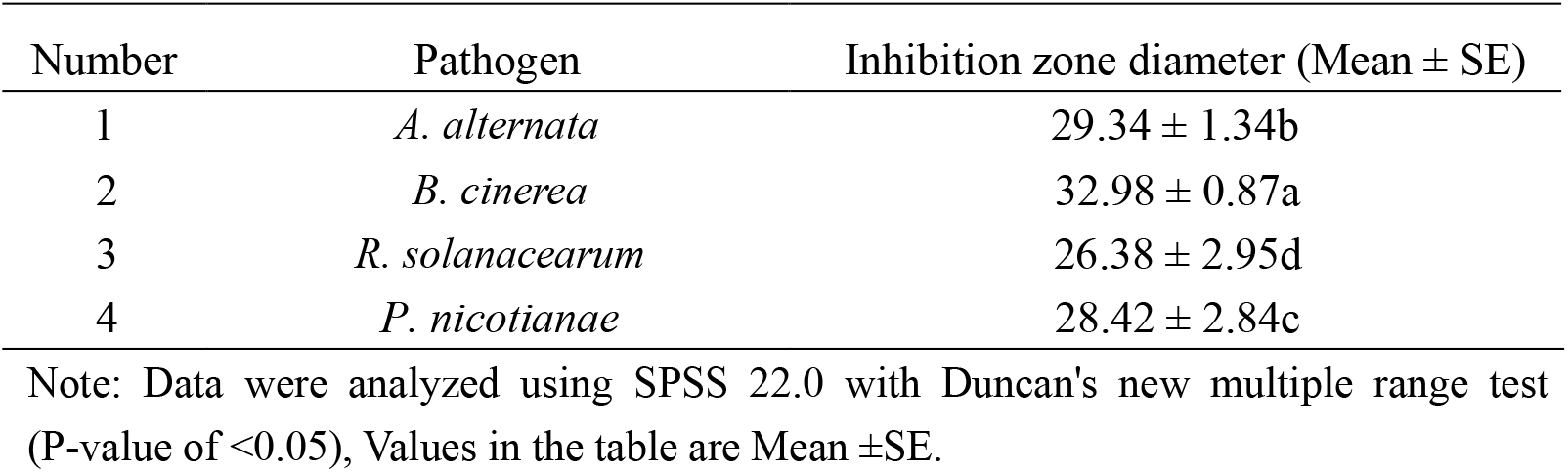
The inhibitory efficiency of strain Ba168 on 4 tobacco pathogens

### MIC of crude extract of strain Ba168 and its effect on the mycelia weight of *P. nicotianae*

The MIC of crude extract of strain Ba168 that completely inhibited the growth of the mycelia of *P. nicotianae* was 5μL/mL. Moreover, it can be seen from Table 4 that mycelia dry weight and wet weight decreased significantly as the concentration of crude extract increased. When the concentration reached 2μL, mycelia dry weight is only 28.3 mg, and the inhibitory effect can be observed apparently. When the concentration exceeded 3μL, mycelia basically stopped growing.

The results suggests that antimicrobial substances produced by Strain Ba168 could strongly inhibited the mycelia growth of *P. nicotianae.*

**TABLE 3.** Effect of crude extract of Ba168 on the weight of mycelia of *P. nicotianae*

| Concentrations of crude extracts of Ba168 (uL/mL) | *P. nicotianae* |  |
| --- | --- | --- |
|  | Wet weight (mg) | Dry weight (mg) |
| 0 | 3610.73 ± 31.94a | 156.00 ± 0.62a |
| 1 | 1366.65 ± 17.76b | 84.45 ± 0.95b |
| 2 | 510.25 ± 3.31c | 28.4 ± 0.34c |
| 3 | 134.325 ± 2.59d | 10.48 ± 0.13d |
| 4 | 99.00 ± 0.56d | 1.73 ± 0.09e |

Note: Data were analyzed using SPSS 22.0 with Duncan's new multiple range test (DMRT) (P-value of <0.05), Values in the table are Mean ±SE.

**TABLE 4.**
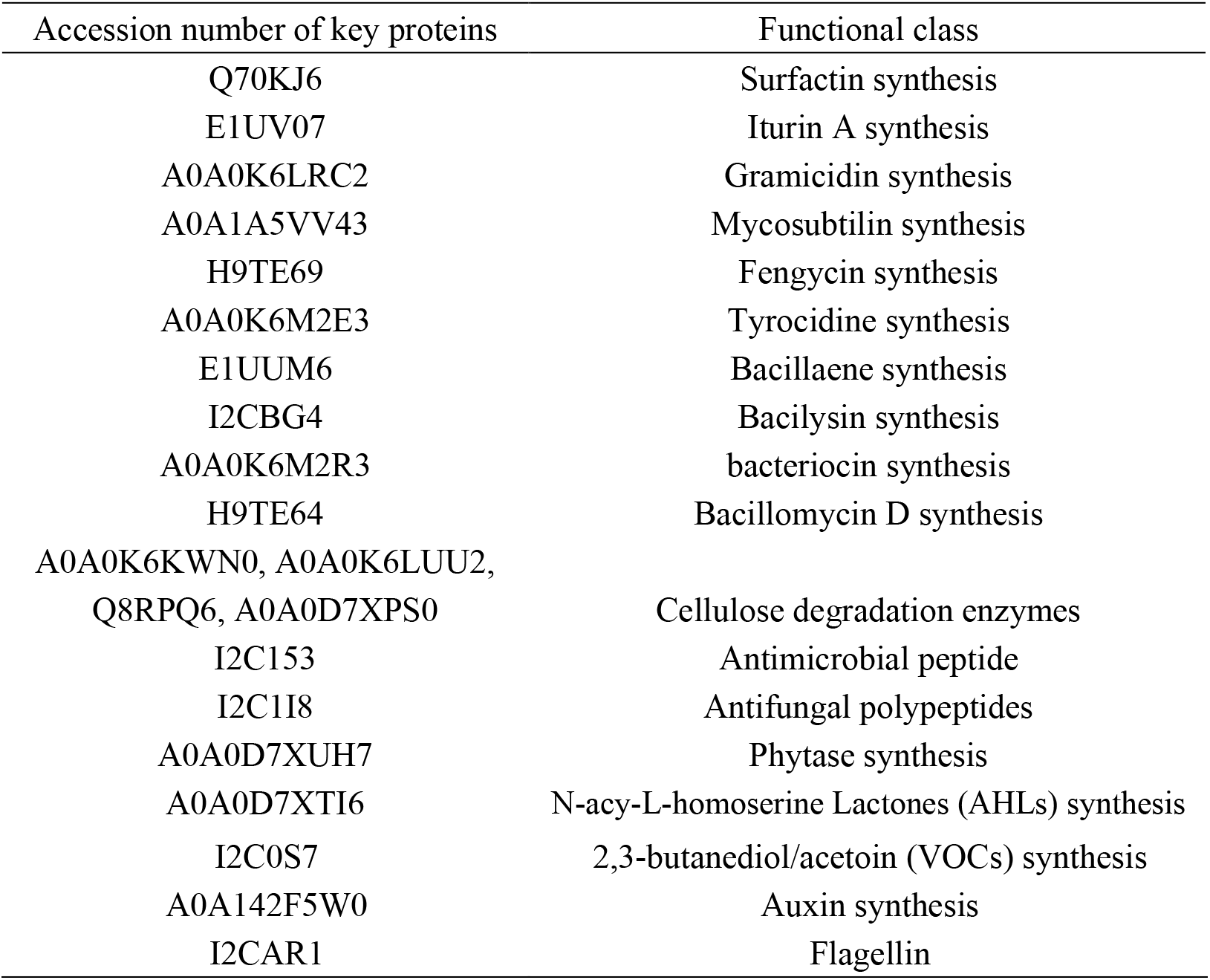
Proteome analysis of crude extract of strain Ba168

### Effects of crude extract of strain Ba168 on Extracellular conductivity and pH of *P. nicotianae*

As can be seen from Fig 2, compared to the control group, the extracellular conductivity of mycelia, exposed to Ba168 crude extract, increased significantly with the extension of treatment time. Moreover, this trend is positively correlated with the concentration of the crude extract, as the concentration of treatment solution increased from 1μL to 4μL, the conductivity values of each group were significantly different. Besides, the conductivity of treatment groups increased continuously over time, and evidently higher than that of the control group. The extracellular conductivity of the four groups reached 94.12 μs/cm, 108.36 μs/cm 116.46 μs/cm 123.13 μs/cm after 120 min, respectively. The effect of crude extract of strain Ba168 on extracellular pH is shown in Fig 3. With the extension of time, the pH showed an overall downward trend. In addition, after the increase of dose, the pH of each treatment group was higher than that of the control group at any time, and the higher the dose, the higher the pH. After 105 min, the concentrations from 0 to 4μl/mL were 5.03, 5.16, 5.31, 5.42 and 5.72, respectively.

**FIG 2.**
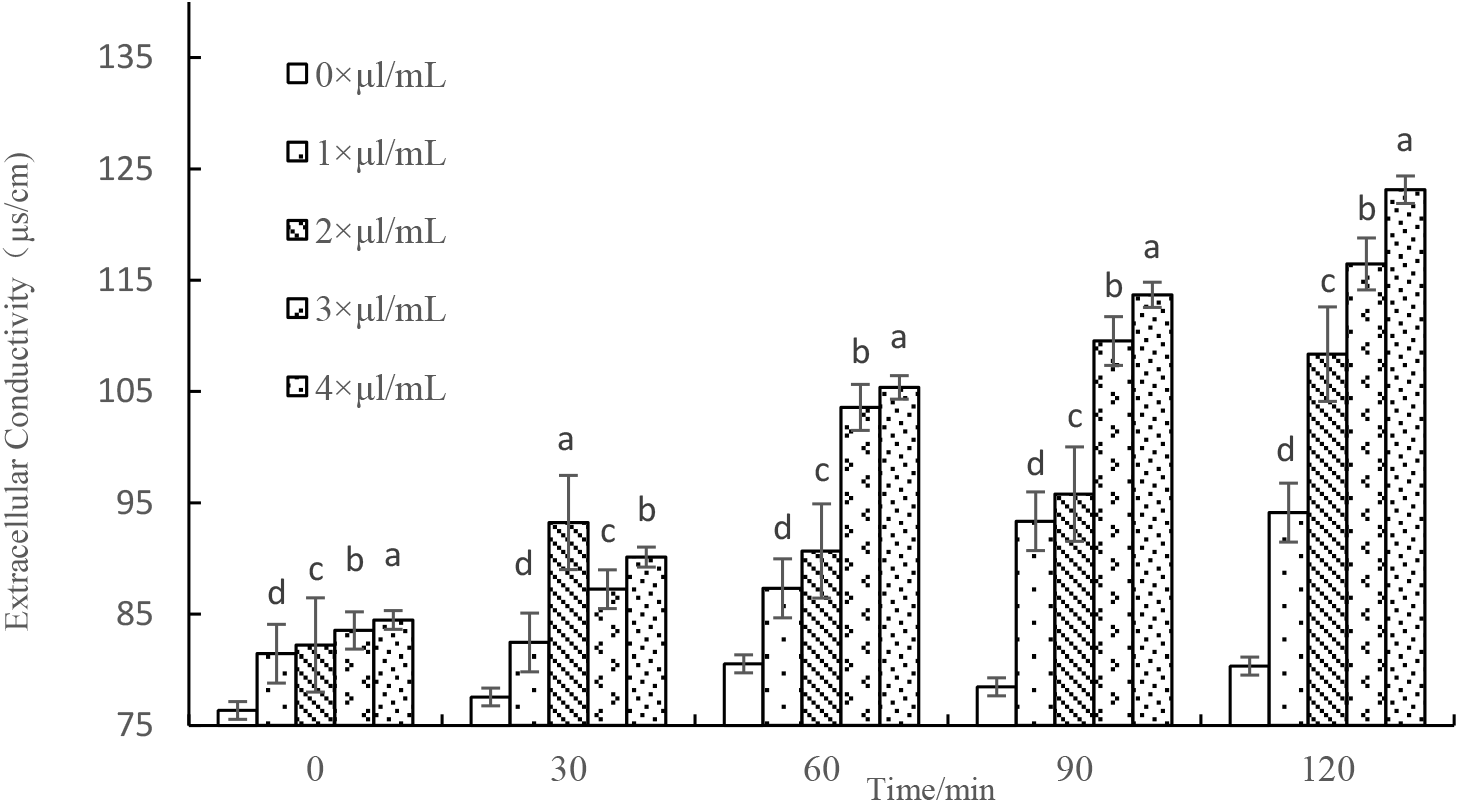
Effect of Ba168 crude extract on extracellular conductivity of *P. nicotianae*

These results indicate that the antifungal activity of Ba168 crude extract may be associated with the irreversible damage to the cytoplasmic membranes and cell wall of *P. nicotianae*, which contributes to intracellular contents leakage from the cells.

### Effect of crude extract of strain Ba168 on the morphology of mycelia of *P. nicotianae*

It can be seen from Fig 4A, 4C that the mycelia of *P. nicotianae* without the treatment of crude extract of strain Ba168 are plump under SEM analysis, and the mycelia’s surface is not damaged or punctured. However, the majority of mycelia presents shrinkage deformation after the treatment of 1 μL/mL Ba168 crude extract (Fig 4B). Interestingly, when we treated the mycelia with the crude extract of 4 μL/mL, as shown in Fig 4D, the mycelia are severely deformed and wrinkled and perforated on the surface of the mycelia.

**FIG 3.**
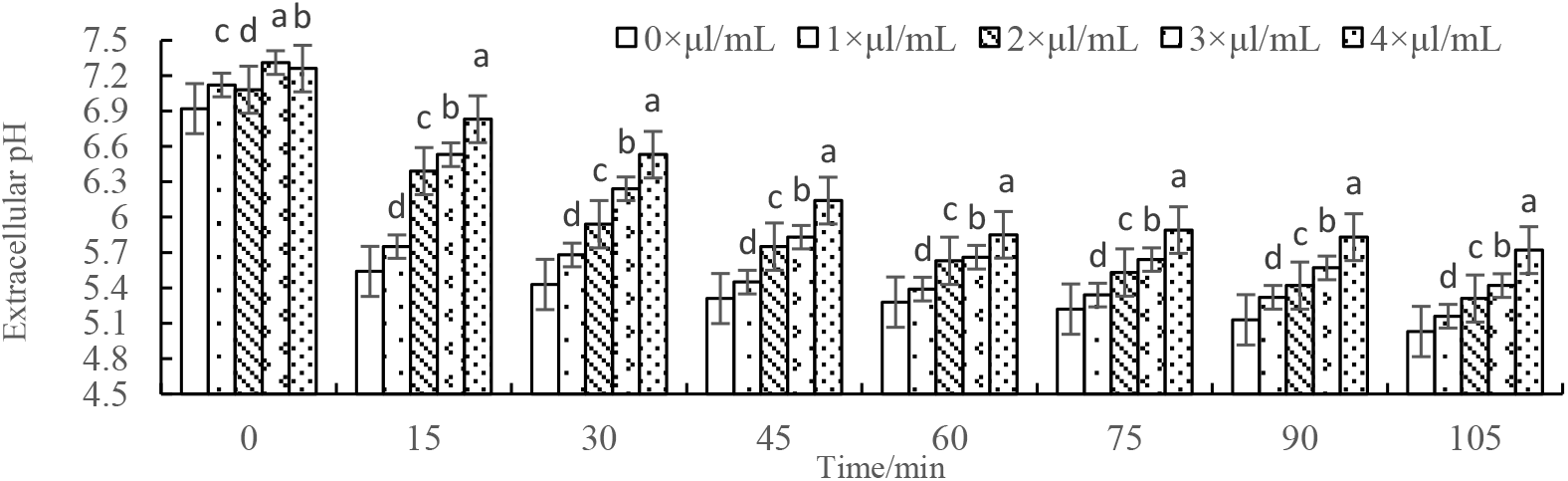
Effect of Ba168 crude extract on extracellular pH of *P. nicotianae*

**FIG 4.**
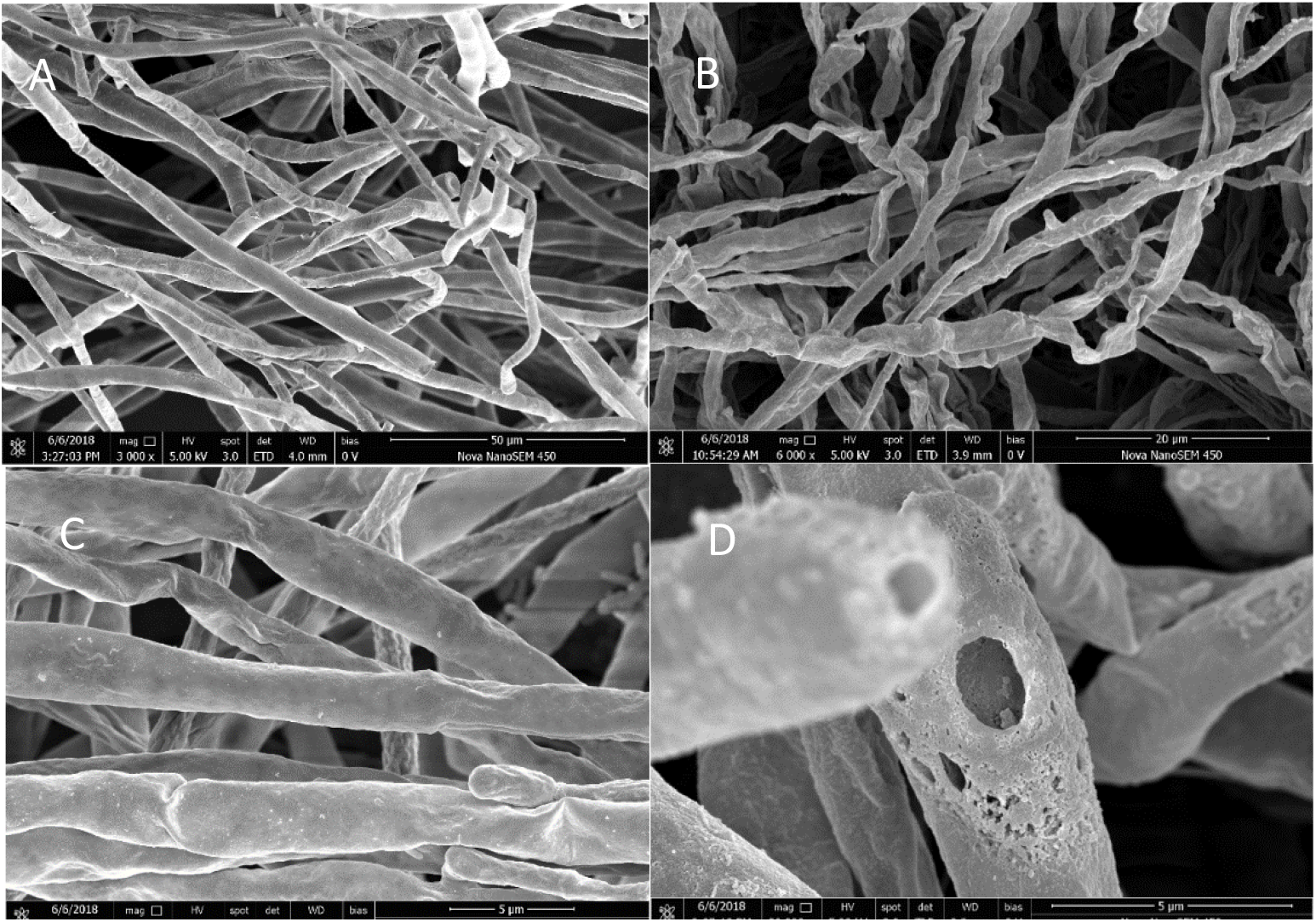
**A** (3000×, bar=50 μm), **4C** (16000×, bar=5 μm): Scanning Electron micrographs(SEM) of the untreated hyphae of *P. nicotianae*. **4B** (6000×, bar=20 μm): SEM of the hyphae treated with 1μl/mL concentration of crude extracts.**4D** (30000×, bar=5 μm): SEM of the hyphae treated with 4μl/mL concentration of crude extracts.

### Proteome analysis for FUP evaluation

Some of the accession number of identified protein shows in Table 4. The identified antifungal polypeptides (I2C1I8), Antimicrobial peptide(I2C153), Cellulose degradation enzymes(CDEs) (A0A0K6KWN0, A0A0K6LUU2, Q8RPQ6, A0A0D7XPS0) may directly responsible for *P. nicotianae* suppression activity *in vitro* studies. Flagellin (I2CAR1) is a classical ISR inducer (49). Phytase (A0A0D7XUH7) help plants to transforming and utilizing insoluble ions in bulk soil and thus have plant growth promoting effect (50). surfactin synthetase (Q70KJ6), Linear gramicidin synthase (A0A0K6LRC2), Iturin A synthetase (E1UV07), Fengycin synthetase (H9TE69), Mycosubtilin synthase (A0A1A5VV43), Tyrocidine synthase (A0A0K6M2E3), Bacillomycin D synthetase (H9TE64), Homoserine kinase (A0A0D7XTI6), Acetoine/ butanediol dehydrogenase (I2C0S7), Tryptophan synthase beta chain (A0A142F5W0), are responsible for Surfactin synthesis, Iturin A synthesis, Gramicidin synthesis, Mycosubtilin synthesis, Fengycin synthesis, Tyrocidine synthesis, Bacillomycin D synthesis, N-acy-L-homoserine, Lactones (AHLs) synthesis, 2,3-butanediol/acetoin (VOCs) synthesis, Auxin synthesis, respectively.

These identified proteins and polypeptides implies that strain Ba168 may have the properties of plant growth stimulation, aggressive colonization and biocontrol.

### Identification of strain Ba168

The pure colonies of strain Ba168 on LB agar medium after 48 h incubation are canary yellow, rough surface, round as a whole with irregular but smooth margin, opaque, drying and no pigment production. The cells of strain Ba168 are rod-shaped, peritrichous, and the spores of it are long elliptic (Observed by scanning electron microscopy (SEM) (Fig 5A) and transmission electron microscopy (TEM) (Fig 5B)). Based on the morphology characteristics described above and the physiological and biochemical characteristics organized in Table 5, as well as the results that received from NCBI database when we using 16SrRNA homology of partial sequences of strain Ba168 to conducting blast, we preliminarily confirmed Ba168 as *B. amyloliquefaciens*. However, the sequence with the highest similarity in blast results without the support of articles is unreliable. We consequently retrieved other represent sequences which published in many articles to conduct the phylogenetic analysis. As is shown in Fig 6, the evolutionary history of Ba168 was inferred by using the Maximum Likelihood method based on the Tamura-Nei model (51).

**FIG 5.**
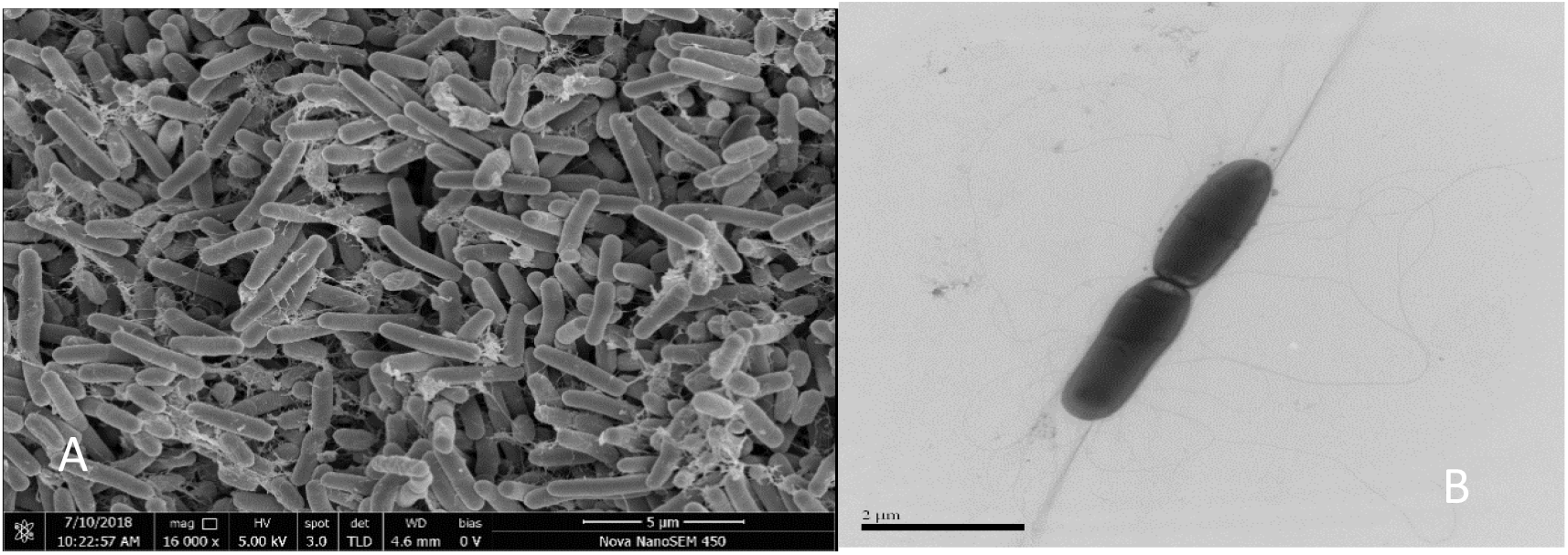
**A**: Scanning Electron micrographs (Nova Nano SEM 450, FEI, American) of spores of strain Ba168. Bar=5μm. **B**: Transmission electron micrographs (HT7700, HITACHI, Japan) of spores of strain Ba168. Bar–2μm

**TABLE 5.**
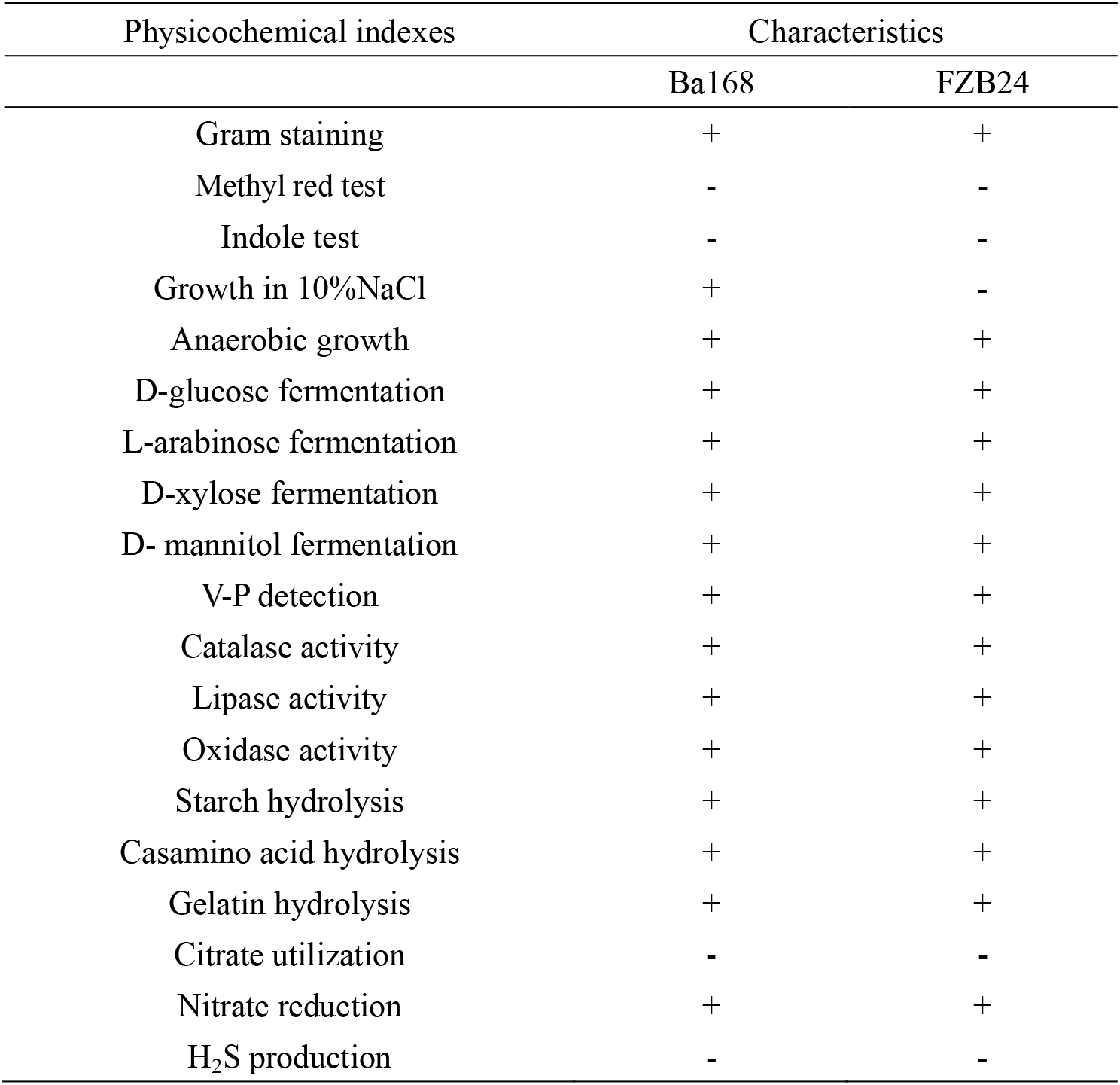
Comparison of biochemical and physiological characteristics of Ba168 strain and *B. amyloliquefaciens* FZB24. Note: “+”Positive; “–”negative.

To summarize, based on morphological, physiological, biochemical, and molecular characterization, strain Ba168 was identified as *B. amyloliquefaciens*.

### Field assays

The results of field trials was demonstrated in Table 6. In Qinyan96, the control efficacy of *B. amyloliquefaciens* Ba168 against TBS is 77.88% ranked behind the *B. methylotrophicus* (83.22%) and MPA (78.38%), but higher than the 80% Dimethomorph water dispersible granules (73.52%), *B. licheniformis* (73.36%), *B. subtilis* (73.98%), *B. laterosporus* (70.46%) and *B. pumilus* (70.37%). The disease control efficacy of Ba168 in NC89 is 66.50 % which is only second to MPA (69.33%) and marginally higher than Dimethomorph (59.60%) and, but significantly higher than *B.licheniformis* (53.58%), *B. subtilis* (56.18%), *B. laterosporus* (54.33%), *B. methylotrophicus* (51.48%), *B. pumilus* (70.37%). After pooling analysis, we found that the disease control efficacy in NC89 is generally lower than in Qinyan 96. Intriguingly, almost all groups decline moderately excepted *B. methylotrophicus* which shows a considerable dip from 83.22% to 51.48%.

**TABLE 6.**
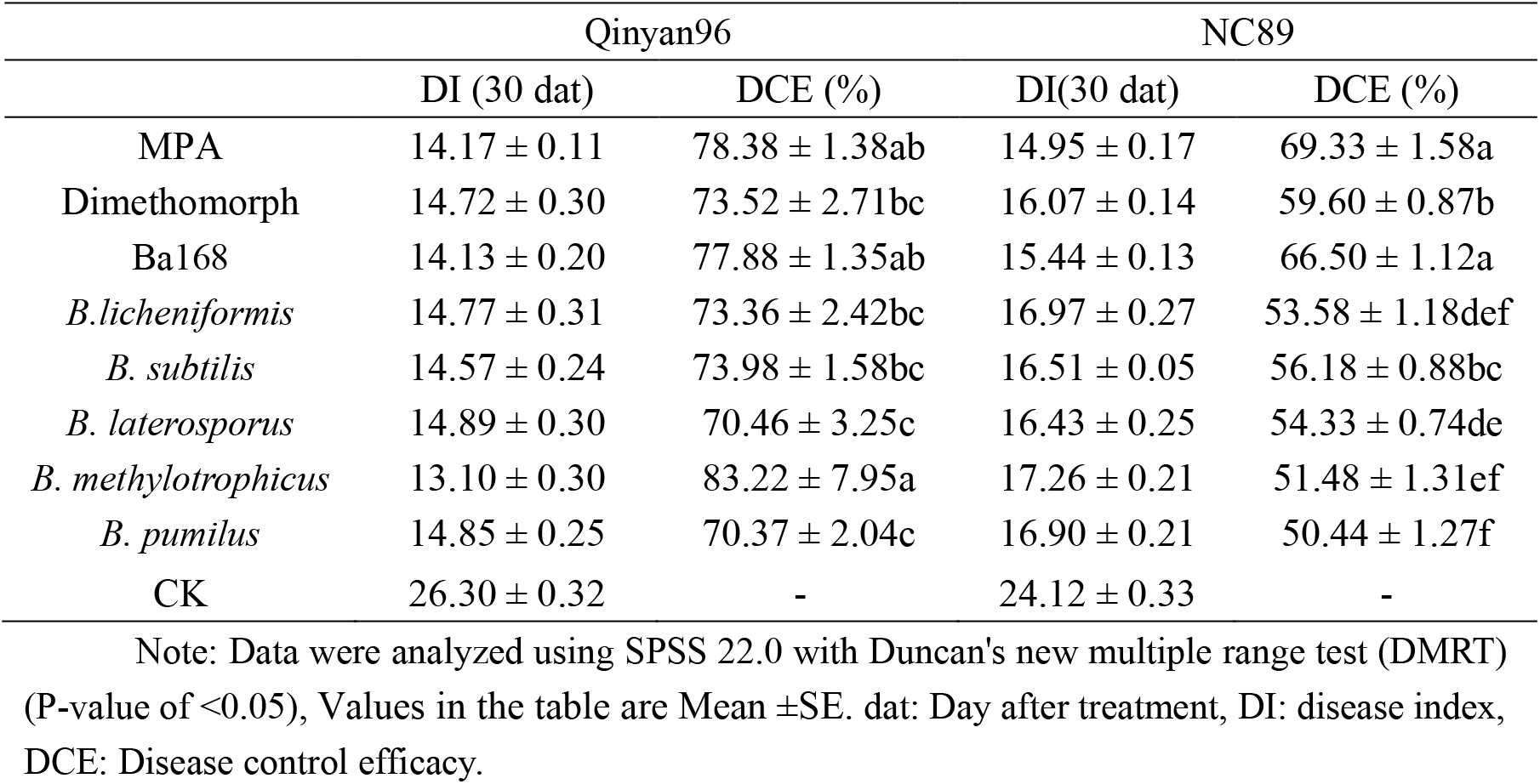
TBS control efficacy in Qinyan96 and BC89 from 2017-2018

## Discussion

In this study, an isolated strain Ba168 was identified as *B. amyloliquefaciens* by its morphology, biochemical and physiological characteristics (Fig 5, Table 5), and 16SrRNA sequence alignment (Fig 6). *B. amyloliquefaciens* has been widely used in food(52), medicine(53), animal husbandry(54), aquaculture(55), agriculture and forestry(56). Although the existence of horizontal gene transfer(57, 58), the ability to be used as a biocontrol strain is possessed by nearly all of the reported strains (59).

**FIG 6.**
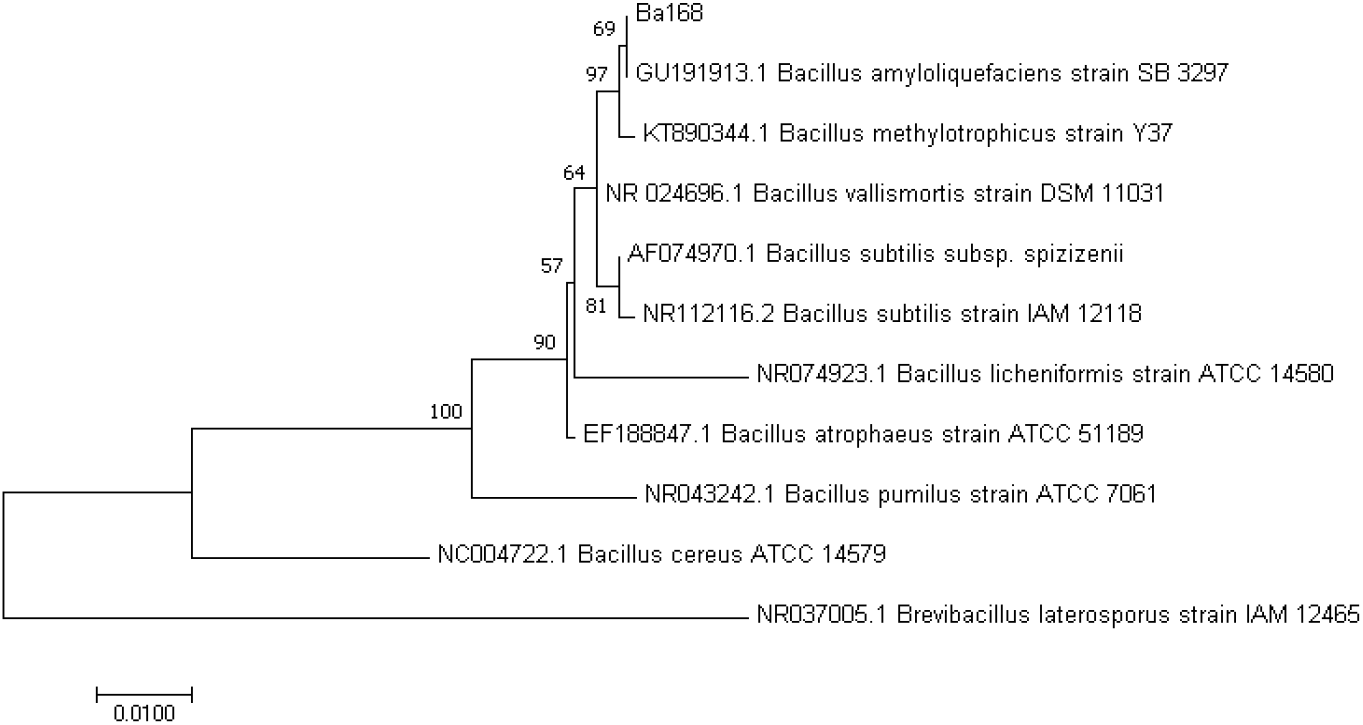
Phylogenetic tree of strains Ba168 base on Maximum Likelihood method analysis of 16SrRNA sequences data. Bootstrap value <51% are not shown. *B. laterosporus* strain IAM 12465 represent the out-group.

*In vitro* studies, *B. amyloliquefaciens* Ba168 was able to strongly inhibit the development of dominant tobacco pathogens and have a better *P. nicotianae* suppression activity than other tested biocontrol strains (Table 1, 2). Extracellular metabolites of a biocontrol strain, to some extent, reflect the biocontrol potential of PGPR after rhizosphere colonization in the field (29). Thus we further focused on the characteristics of the crude extract of culture filtrates of Ba168. The crude extract of it present a strong antifungal activity (MIC = 5 μL/mL) which is more effective than eugenol (35), *Trichoderma* EtOAc extract (13) and mixtures of plant extract (60). Mycelia growth can be severely inhibited when exposed to Ba168 crude extract (Table 3). Intriguingly, the data of extracellular conductivity (Fig 2) and extracellular pH (Fig 3) shows that Ba168 crude extract causes the outflow of mycelium contents and in a dose-dependent way. We are not surprising that the crude extract of strain Ba168 could exert irreversible damage on the cell membrane of *P. nicotianae* and ion channels on the membrane. That is reminiscent of the function of surfactin, Iturin, and fengycins families, for which surfactin families can easily combine with the lipid layer of the cell membrane to damage the integrity of the membrane (61); the iturin class does not destroy the membrane structure, but it can increase the membrane permeability and form ionic conduction hole to interfere with the transport of transmembrane substances (62); the mechanism of fengycins is unclear, but they also tend to interact with lipid layers and retain the potential for dose-dependent changes in cell membrane structure and permeability (63). This makes the recorded SEM reasonable in Fig 4B, in which the mycelia wrinkles severely. Theoretically, these lipopeptides should present in the crude extract under this sample preparation method, but the molecular weight of them identically lower than 5000, so they were not identified in this study. Whether these antibiotics play a role in this study need further vindication.

When the concentration of Ba168 crude extract comes to 4μL/mL, mycelia cell walls are dissolved (Fig 4D) to a certain extent. Interestingly, as the cell walls of *P. nicotianae* are composed of cellulose (64), we can infer that CDEs are also involved in Ba168 crude extract. In proteome assays, CDEs including beta-glucanase, endoglucanase, pectin lyase and pectate lyase, were identified. CDEs either from plants or microbial organisms may also play a role in anti-oomycete activity as reported in recent studies (65). For instance, beta-1,3-glucanase from pepper and tomato can inhibit hyphal growth of *Phytophthora capsici* (66) and cellulases from *Bacillus* are effective in degrading the cell wall of *Phytophthora* pathogens (65). Nevertheless, extensive cell wall digestion of *P. nicotianae* may do not bring death to oospores or a mycelial piece, since new hyphae grew even if others were being actively digested (67, 68). Besides, some antimicrobial peptides(69) and antifungal polypeptides (70) were also detected in Ba168 crude extract. Therefore, it is speculated that *P. nicotianae* suppression activity may be synergized by the substances containing in the crude extract of *B. amyloliquefaciens* Ba168.

In FUP evaluation (Table 4), the identified phytase (50), flagellin (49) and other proteins related to the synthesis pathway of antibiotics(like surfactin (71), tyrocidine (72), gramicidin (73), bacillomycin D(74), Bacillaene, Bacilysin, bacteriocin (70)), auxin (75), N-acy-L-homoserine Lactones (AHLs) (76), 2-3-butanediol (VOCs) (77), implies the comprehensive biological control capability of strain Ba168 including aggressive colonization (9, 23), induced systemic resistance of host plants (19) and plant growth-promoting (78). In general, lab studies showed that *B. amyloliquefaciens* Strain Ba168 have a good *P. nicotianae* suppression activity and FUP. However, whether it works under field conditions remains unknown. And, to our knowledge, there is no available literature about the biocontrol efficacy of *Bacillus* strains against TBS caused by *P. nicotianae* under field.

TBS is notoriously hard to control. The plant protection toolbox for this soil-borne disease remains undeveloped. In field studies, we provide a complete set of techniques for the application of PGPRs in field control of TBS including how to prepare bioagents and how to use them in field. The combination application method similar to K. Elanchezhiyan et al(79) was adopted to evaluate the TBS suppression activity of strain Ba168 and its rivals. Results show that the disease control efficacy of strain Ba168 is lower than MPA in two tobacco varieties, but higher than Dimethomorph preparations. Dimethomorph, Propamocarb, Azoxystrobin all have been applied in TBS control for many years, drug-resistant oomycetes emerge (80). Despite that, Dimethomorph is more effective than Propamocarb, Azoxystrobin when separately used (80). But, in this study, we show that Mixture of Propamocarb and Azoxystrobin is better than Dimethomorph used alone. *B. amyloliquefaciens* Ba168 is more favorable in aspect of biocontrol stability and suppression activity in TBS control, compared to *B. licheniformis, B. subtilis, B. laterosporus, B. methylotrophicus, B. pumilus.* Interestingly, the disease control efficacy under field conditions conducted in Qinyan96 is generally higher than that in NC89. Since all the experiments were conducted in the same sites and the climate difference was marginal. It is suggested that QingYan96 is more suitable than NC89 to be planted as a cultivar which resistant to TBS in southern Shaanxi. Additionally, the relative disease control efficacy of *B. methylotrophicus* against TBS had the greatest drop in two varieties, from (83.22 ± 7.95)% to (51.48 ± 1.31)%. This may because *B. methylotrophicus* are methylotrophy (81, 82), which are more restrict in application conditions.

In conclusion, results from our study indicate that *B. amyloliquefaciens* Ba168 has the capability to be utilized in the biocontrol of TBS under field conditions. Besides, although the differences exists in TBS control efficacy, the strategy of PGPR application is promising in this soil-borne disease control. While the disease control efficacy of strain Ba168 is slightly lower than that of MPA, it is environmentally friendly and will generate no residues after applied many years. However, in order to expand the application prospect of strain Ba168, further studies need to focus on the research of mixed microbial agents (83) or mixtures of biocontrol agents and harmless pesticides, containing strain Ba168. If possible, simplified the application method without reducing control efficacy is also on our agenda. On the other hand, it requires a combination of ingenuity and hard work to finding an effective biocontrol PGPR for practical applications(5). In this study, the proteome analysis was used as a supplementary data to evaluated FUP of strain Ba168. With the decrease of the cost of transcriptome and proteome tools, these tools may help us to simplify PGPR-finding process.

## ACKNOWLEGEMENTS

For providing us with the scanning electron microscope (SEM), transmission electron microscopy (TEM) and proteome analysis we thank Life Science Research Core Services (LSRCS) and The State Key Laboratory of Crop Stress Biology for Arid Areas. For the field trials, we thank Fu Qiang. For the language revision of this article, we thank Elizabeth, Sun Guangzheng. For the identification of strain Ba168, we thank Feng Zhizhen. For the experiments about extracellular conductivity and pH, we thank Wang Lei and Zhu Yunqi. For revision advises we thank Professor Sun Guangyun and Wang Yang.

